# Applying And Promoting Open Science In Ecology - Surveyed Drivers And Challenges

**DOI:** 10.1101/2021.10.12.464125

**Authors:** Christian B. Strømme, A. Kelly Lane, Aud H. Halbritter, Elizabeth Law, Chloe R. Nater, Erlend B. Nilsen, Grace D. Boutouli, Dagmar D. Egelkraut, Richard J. Telford, Vigdis Vandvik, Sehoya H. Cotner

**Affiliations:** Department of Biological Sciences, University of Bergen, Thormøhlensgate 53A, Bergen, 5006, Norway; Norwegian Institute for Nature Research, Høgskoleringen 9, Trondheim, 7034, Norway; Department of Biology Teaching and Learning, University of Minnesota, Minneapolis, Minnesota, 55108, USA; Working Conservation Consulting, Fernie, BC, Canada

## Abstract

Open Science (OS) comprises a variety of practices and principles that are broadly intended to improve the quality and transparency of research, and the concept is gaining traction. Since OS has multiple facets and still lacks a unifying definition, it may be interpreted quite differently among practitioners. Moreover, successfully implementing OS broadly throughout science requires a better understanding of the conditions that facilitate or hinder OS engagement, and in particular, how practitioners learn OS in the first place. We addressed these issues by surveying OS practitioners that attended a workshop hosted by the Living Norway Ecological Data Network in 2020. The survey contained scaled-response and open-ended questions, allowing for a mixed-methods approach. Out of 128 registered participants we obtained survey responses from 60 individuals. Responses indicated usage and sharing of data and code, as well as open access publications, as the OS aspects most frequently engaged with. Men and those affiliated with academic institutions reported more frequent engagement with OS than women and those with other affiliations. When it came to learning OS practices, only a minority of respondents reported having encountered OS in their own formal education. Consistent with this, a majority of respondents viewed OS as less important in their teaching than in their research and supervision. Even so, many of the respondents’ suggestions for what would help or hinder individual OS engagement included more knowledge, guidelines, resource availability and social and structural support; indicating that formal instruction can facilitate individual OS engagement. We suggest that the time is ripe to incorporate OS in teaching and learning, as this can yield substantial benefits to OS practitioners, student learning, and ultimately, the objectives advanced by the OS movement.

## Introduction

Open Science (OS) encompasses a broad set of principles and practices that are aimed at generally improving the accessibility and transparency of scientific research [1]. Although a unifying definition has yet to emerge, OS can be understood as concurrent trends in science and society broadly aiming to enhance openness about and free and inclusive access to all aspects of science, driven mainly by networks of individual OS practitioners [2,3]. Different facets of OS, each emerging from slightly different assumptions and goals, focus on promoting diverse and equitable access and contribution to knowledge; innovation and efficiency through collaboration; quality and credibility through transparency; access to open research platforms, new and efficient tools and services; and alternative metrics for assessing research contribution and impact [3,4]. Together, the rapidly evolving principles and practices associated with OS are expected to revolutionise how research is done and shared in the not-too-distant future [5].

The transition towards OS has required the development of necessary infrastructure, including platforms for collaboration and large-scale interactive databases, and is currently re-defining publishing models (e.g. Plan S, https://www.coalition-s.org). This transition is supported by developments in licencing, data and metadata standards, and by requirements for open publishing and data sharing set by research funding bodies. While many practitioners adopt a subset of OS principles and practices for idealistic or pragmatic purposes, institutional support has been crucial for developing OS infrastructure and mainstreaming OS. Moreover, OS principles conceptualised by practitioners can be adopted and implemented by institutions, as exemplified by the inclusion of the FAIR guiding principles [6] by the European Open Science Cloud [7] and the newly adopted UNESCO Recommendation on Open Science [8]. Through such efforts, many of the key institutional, economic and infrastructure-related challenges in the transition to Open Science have been addressed.

Despite these developments, the practices associated with OS are not widely implemented across research communities, but rather within select groups, networks or events involving OS practitioners, or in a sub-optimal piecemeal manner [9]. Thus, a major challenge for fully utilising the potential of OS involves its widespread uptake by diverse members of the scientific community, which will require a major cultural and behavioural shift among scientists. Such a transition is not straightforward, considering the variability in how individual researchers perceive OS in terms of values, required skills and pragmatic trade-offs between benefits and costs [10]. For some practitioners, the interest and entry point to adopting OS practices may be driven by the necessity of reproducibility and replicability. In addition, parallel networks of researchers can differ in their emphasis of these and other OS aspects as well as the extent of collaboration among peers [4]. Therefore, considerable variability in how OS is understood and practiced can be found both between individual researchers and between groups or networks.

In the light of these challenges, an understanding of how OS practices and principles are learned, understood, and transmitted among researchers can better inform institutions and policymakers invested in implementing OS in full. As the OS movement is still relatively young, we are particularly interested in practitioners’ thoughts on the role of OS in teaching and supervision. Arguably, for OS to become an integral part of mainstream science, its inclusion in how science is taught and learned may prove highly effective and necessary. To characterise the variation in how OS is used, understood, and perpetuated in a network of OS practitioners, we surveyed attendees at an international two-day workshop dedicated to openness and transparency in applied ecology, organised by the Living Norway Ecological Data Network (livingnorway.no). This peer-driven collaborative initiative was established in 2019 with the purpose of improving management of ecological data from Norwegian research institutions, being closely associated with the Norwegian participant node in the Global Biodiversity Information Facility (GBIF, https://www.gbif.org).

Attendees at the 2020 international Living Norway Colloquium were invited to answer a digital survey developed to address the following research questions:

- How is OS perceived among practitioners in ecology?
- Which OS aspects do practitioners interact with, and how frequently?
- What are the perceived benefits and risks for individual engagement in OS?
- Which OS aspects have practitioners encountered in their own formal education?
- How do OS practitioners involved in higher education value OS in teaching and supervision of their students?

## Methods

### Colloquium

The 2nd International Living Norway Colloquium was a hybrid event in October 2021 where participants could either attend in person or join via a digital meeting platform. For two days, attendees participated in thematic sessions consisting of plenary talks, plenary and group discussions, and group assignments (see Supporting information S1 Table for the full program).

### Survey development and administration

Our investigative team included individuals involved in organising the colloquium itself and associated collaborators. We met twice before the colloquium to clarify research questions, develop survey items, and subject the items to talk-aloud refinement. This resulted in a questionnaire that we structured into three parts where each part was distributed to colloquium participants as follows: Part I three days before the event started, Part II at the end of the first day, and Part III after the second and final day of the event. We split the survey partially to distribute the effort of respondents taking the survey, to focus on different themes, and to gather data on whether participants’ understanding of OS evoloved during the event.

### Survey structure and content

The questionnaire consisted of a combination of Likert-scale, constrained choice, and open-ended questions. Constrained choice questions were used to obtain background information such as degree, affiliation, gender, and experience with OS aspects. Likert-scale response questions were related to experience with and perceived importance of different OS aspects to research, teaching, and supervision of students engaged in thesis work. Open-ended questions asked colloquium participants to define OS and describe what has helped and hindered their OS engagement. To link the submitted responses for each of the three parts, survey participants were asked to provide their email as an identifier with the assurance that this identifier would be removed within two weeks of the event. For the complete survey questionnaire, see Supporting information S2&3 Tables.

### Data management

We uploaded and distributed the electronic survey using SurveyXact (Ramboll, Denmark), and survey responses were submitted and collected through this provider. Email addresses submitted by respondents were deleted within two weeks and remaining data were uploaded to the GitHub repository together with the R code. We clarified management procedures for personal information with NSD (Norwegian centre for research data).

### Qualitative analysis

We established a team (GDB, SHC, and AKL) that coded the open-ended responses to the following three survey items: definitions of OS, what hinders-, and what helps individual OS engagement (see Supporting Information S2&3 Tables for the full questions).

Specifically, the team subjected item responses to iterative rounds of inductive coding, beginning with a meeting to establish an initial codebook by reading all responses to a particular question and identifying common ideas [11]. Because the open-ended questions served as opportunities to elicit new ideas from this novel participant group, we did not begin with *a priori* categories and rather remained open to all ideas that were present in the data. For each question, after describing initial categories, at least two of the three coders independently coded all responses. The coders then met and discussed differences along with any additional ideas that did not fit into the original categories. The coders repeated this process until no new categories were needed to capture all relevant ideas. Final codebooks can be found in Supporting Information (S4;S7-8 Tables). For the final coding step, at least two coders coded all responses an additional time using the final codebook and then met and came to consensus on the coding for all responses. Importantly, all coding was done before any coders were aware of the results of the quantitative analyses (see below), to protect the integrity of the coding process. We then discussed overarching themes that resulted from the coding analysis during writing of the manuscript. We have lightly edited some of the quotes reported for grammar and clarity.

### Quantitative analysis

Based on the study aims, we formulated a set of predictions that were preregistered in order to avoid post-hoc hypothesising (see file containing preregistered predictions on the GitHub project repository (https://github.com/christianstromme/LivingNorway2020). We tested the following preregistered predictions:

a. Researchers with academic affiliation have engaged more in open science-related practices compared to other researchers, and more so for early-career researchers.
b. Colloquium participants use open data more frequently rather than contributing with open data and code.
c. Colloquium participants are more likely to use OS in supervision and teaching when they use it in their own research.
d. Colloquium participants that have been taught open science-related practices are more likely to engage in those practices.
e. Colloquium participants perceive OS practices to be more important in their research compared to teaching and supervision.

We performed all quantitative analyses using R software (R Core Team 2020) and the code is available on the GitHub project repository.

We conducted analyses of quantitative data as follows. We treated the scaled responses as ordinal data in our analyses by using cumulative link models (clms, function *clm*) or cumulative link mixed models (clmms, function *clmm*) in the R package Ordinal [12].). For determining the global model structure, namely which fixed terms (including interactions) to include, we followed the pre-registered predictions (see above). Some of the preregistered predictions (numbered in the file on GitHub) yielded overlapping model structures (predictions 1.2 and 3.1), while some predictions could not be tested due to insufficient data (prediction 1.4). Therefore, some of the preregistered predictions were not included further in the study and the specific reasons are stated at the top of each test in the code for quantitative analyses (accessible on GitHub). For analyses that included multiple responses from each survey participant, we included a random term (individual, a numeric labelling variable) in the global model. We determined the structure of the most parsimonious model through a model selection procedure by using the function *dredge* in the MuMIn R package [13]. For all model selection steps, we determined the inclusion of fixed- and random terms based on the lowest AIC estimate. When multiple parsimonious models had a difference of AIC ≤ 2, we selected the model having the simpler structure as the final model. For global- and final model structure, see Supporting Information S5-6 & S9-10 Tables.

For both clms and clmms, final models were checked for violation of the proportional odds assumption, namely if any of the fixed term estimates varied with the response categories. As these fixed terms were nominal factors, nominal tests were conducted by using the function *nominal_test* for clms. As the same function does not apply to clmms, we performed nominal tests for these models through likelihood ratio tests, comparing the most parsimonious model and a model with the same structure except having the fixed factor in question specified as nominal in the formula, thus relaxing the proportional odds assumption. We considered nominal effects significant when model comparisons in the likelihood ration tests yielded *P*<0.05. If these tests revealed violations of the proportional odds assumption, we relaxed this assumption in the final model using the term nominal for the fixed term in question. This setting allowed the regression parameters to vary between different levels of a given covariate for which the proportional odds assumption was violated.

## Results

### Surveyed respondents

Among 128 registered participants, 60 participants completed Part I, 51 completed Part II and 38 completed Part III. Four colloquium participants that responded to Parts II and III did not respond to Part I. The majority of surveyed colloquium participants were affiliated with universities (N=37), followed by research institutes (N=22), other affiliations (N=5) and governmental agencies (N=2). Among these, six respondents stated dual affiliations. Further, the majority of respondents stated Norway as their country of work or study (N=45), followed by countries outside the EU (N=13) and within the EU (N=8) (Supporting information S1-2 Figures). Among these, four respondents stated multiple countries of affiliation. In terms of gender, 27 participants identified as women, 32 as men, and 1 as non-binary. In statistical analyses where Gender was used as a fixed term, the latter category was omitted as a factor level as it was represented by a single observation.

### How do respondents define open science?

We asked colloquium participants to respond to the following open-ended question:

*People define ‘Open Science’ in many ways, and it is a multi-faceted concept. We are interested in how you define Open Science, especially as it pertains to your own work*.

Several codes emerged from the participant responses (Supporting Information S4 Table). Quantifying the occurrence of these codes allows us to infer shared, as well as less common and less salient, perceptions of the meaning of OS (Table 1). In the pre-event responses, some patterns are evident, in that *shared data* was the most frequently identified code in the volunteered definitions of OS, with the vast majority (50 out of 60 respondents) indicating that in OS, data are available for anyone to use (e.g., “Sharing published data and/or raw data in open repositories”). The other most frequent statements included *data availability/accessibility* (38), *sharing codes/methods* (36), *transparency* (28) or *open access publications* (24) (Table 1). For example, one respondent commented, “open science means open access to scientific publications and open sharing of data and code for analysis for scientific publications” (coded as *sharing codes/methods, sharing data*, and *open access publications*). Another shared that “Open science is transparent and repeatable. In ecology there is a particular need for data sharing, i.e. giving colleagues access to raw data for repeating analyses and/or applying alternative methods to extract information from the empirical data” (*transparency, sharing data, replication/reproducibility*).

**Table 1.**
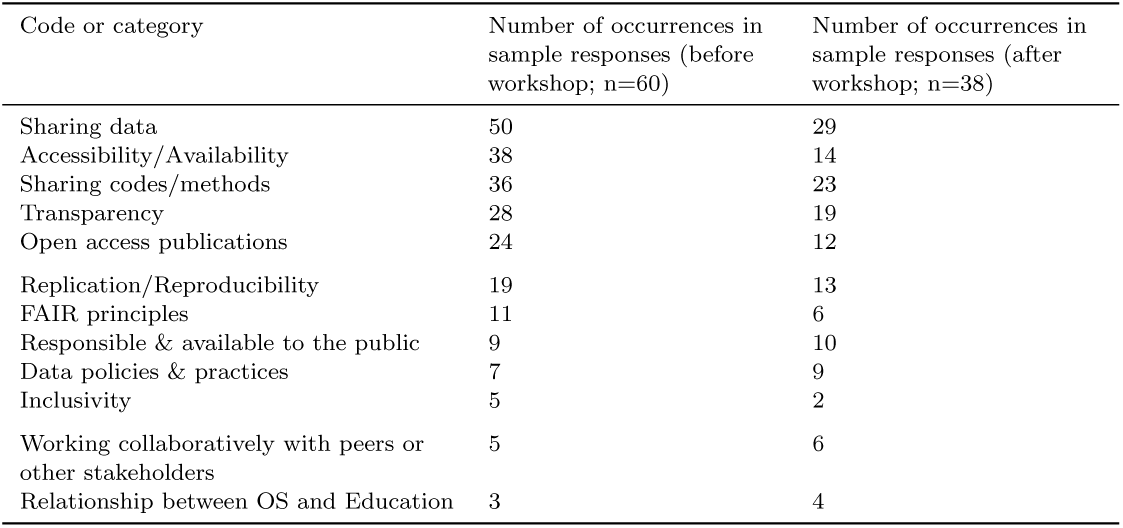
Frequency with which emergent codes were identified in the survey responses in Parts I and II. Codes are organized from most frequent (“sharing data”) to least frequent (“relationship between OS and education”), in Part I.

Very few (3 of 60 respondents) defined OS in a way that specifically referenced education, or a relationship between OS and education, suggesting that it was not as salient of an idea among this population. In all three of these cases, education was mentioned as one of many facets of OS. For example:

“Open science is the ideal of free and accessible availability for everyone to all components of the scientific cycle. Open science entails open education, open research protocols, open methodologies, open data, open code, open data management and analysis, open research publication opportunities, open research readership opportunities, open data synthesis, open science-policy interface, and an open research funding and open science system, science policy and science management.”

Similarly, few responses included inclusivity in their definitions. Those that did reference inclusivity described it as an important reason for OS. For example, one respondent shared:

“I would define Open Science, in the immediate sense, as a way of conducting research in a manner that is transparent (i.e. showcasing/sharing how you are doing your research), however this should extend past just sharing your research and it should also include creating and cultivating a research environment that is open and inclusive to all. Within my work the aspect of reproducibility (particularly code-sharing) is something that I place particular value on.”

### Which OS aspects do practitioners interact with, and how frequently?

Among the 60 respondents to Part I of the survey, 49 were engaged in primary research, and for these we found strong evidence that those having an academic affiliation interacted with OS practices more frequently than colloquium participants with other affiliations (*P* = 0.003) (Fig. 2, Supporting Information S5 Table). Results also indicated that the frequency of interaction with OS practices in general among respondents was higher for men than for women (*P* = 0.004) (Supporting Information S5 Table). However, we did not find evidence for higher OS engagement among early-carreer researchers, as the term was discarded in the model selection process (Suppoprting Information S5 Table).

Respondents stated *Read open access publications* as the OS aspect that was most frequently engaged with, followed by *Used open code* and *Used open data* (Fig. 1). Further, we found strong evidence for respondents using open data and code more frequently than sharing data and code (*P <* 0.001), and men did so more frequently than women (*P* = 0.002) (Supporting information S5 table).

**Figure 1.**
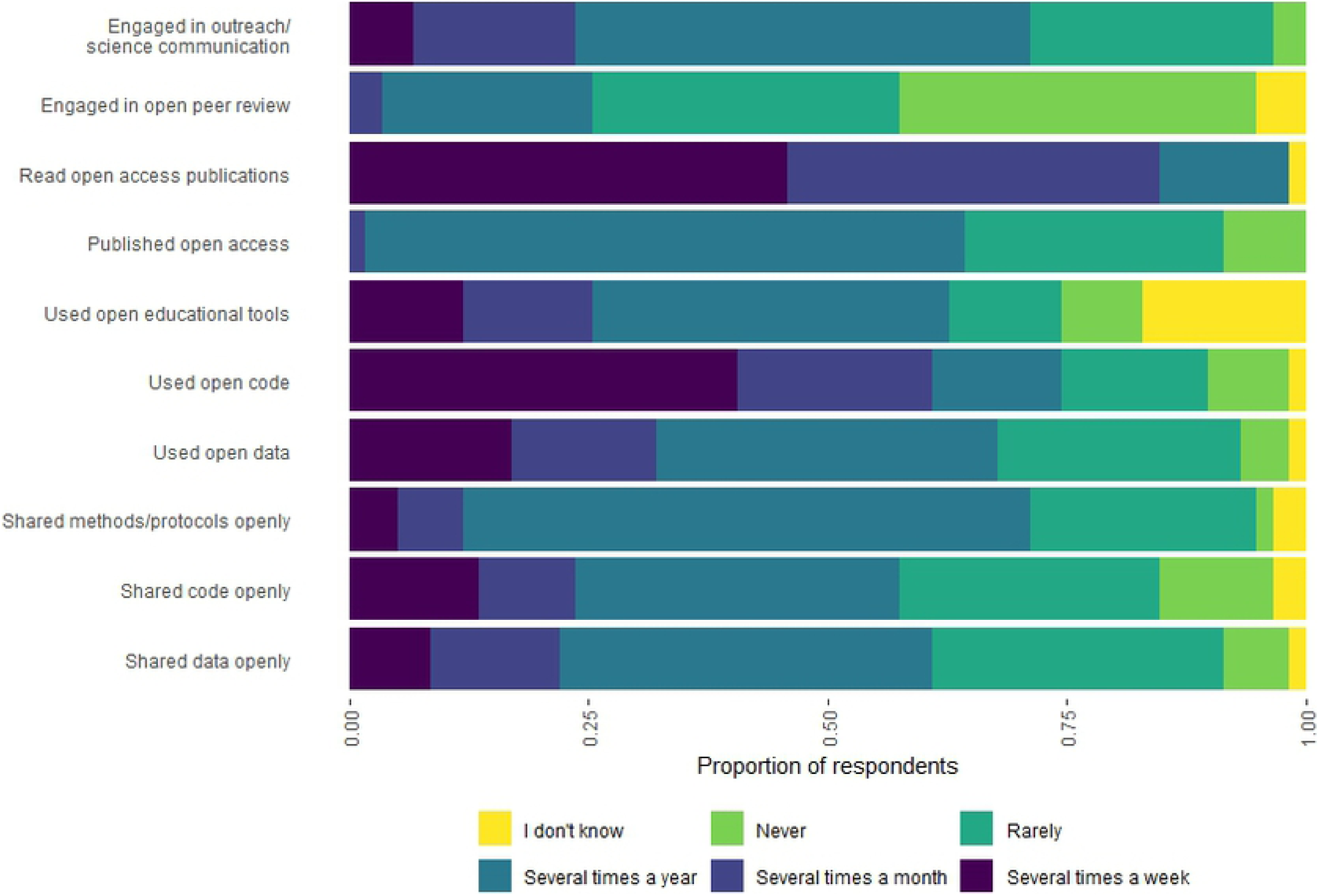
Colloquium participants’ stated frequency of engagement with OS practices.

**Figure 2.**
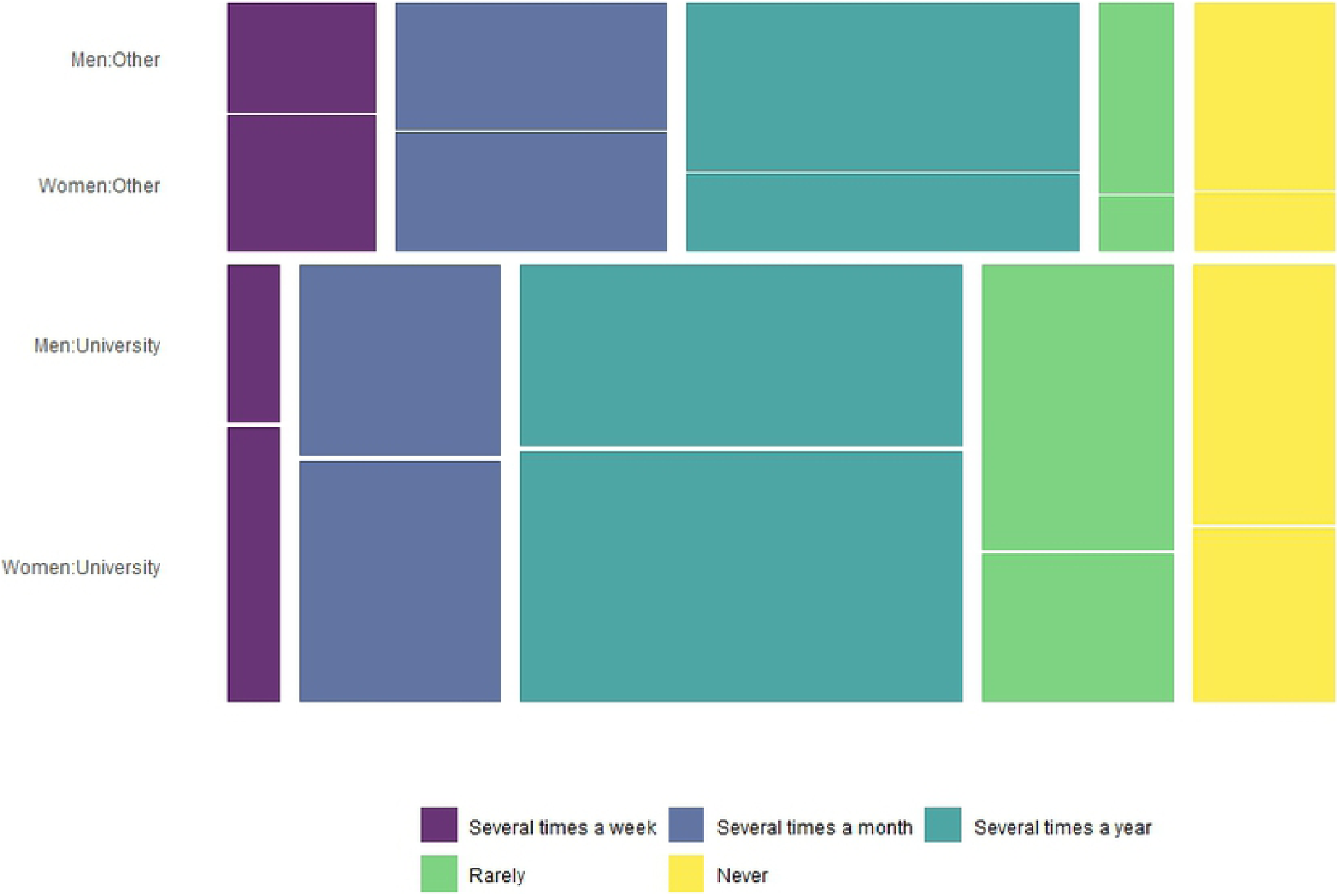
Frequency of engagement with Open Science aspects for surveyed researchers (N=36) participating at the colloquium in relation to affiliation and gender. Height of tiles corresponds to the number of participants for the respective categories of participants, width of tiles corresponds to frequency of scale category.

### What are the perceived benefits and risks for individual engagement in OS?

Participants were asked, via open-ended questions, to share what has hindered them from engaging in OS, and what has helped them to engage in OS. We developed separate codebooks for the “hinders” (Supporting information S7 Table) and “helps” (Supporting information S8 Table) responses. It is important to note that many of the perceived hinders reflect the opposite of what is perceived to help and vice versa.

Of the sixty responses to the “hinders” item, 7 colloquium participants wrote either *nothing* or *N/A*. Many respondents reported *lack of guidelines* (n=15), *lack of time* (15), or *insufficient knowledge* (15) as barriers for engaging in OS (Table 2). One respondent exemplifies these sentiments (along with fear of critique) with the following: “lack of familiarity with relevant online platforms, software, methods… perception that the landscape of the above tools changes very quickly, and keeping up is a big time commitment… fear of doing it wrong.”

**Table 2.** Frequency with which emergent codes were identified for the responses to “what hinders your engagement in OS?” prompt. Codes are organized from most frequent (“Lack of guidelines”) to least frequent (“Fear of critique”)

| Code | Number of occurrences in sample responses (Part I; n=60) |
| --- | --- |
| Lack of guidelines | 15 |
| Time | 15 |
| Insufficient knowledge | 15 |
| Collaborators not using OS | 13 |
| Other/Vague | 12 |
| Cost | 11 |
| Nothing | 7 |
| More work | 5 |
| Insufficient incentives | 5 |
| Legal concerns | 4 |
| Want to get credit | 3 |
| Fear of critique | 3 |

Of the sixty responses to the “help” item, 20 participants referenced *social support* as something that helped them to engage in OS (Table 3). Twenty people also cited *resource availability*. One respondent covered both of these codes saying:

**Table 3.** Frequency with which emergent codes were identified for the responses to “what helps your engagement in OS?” prompt. Codes are organized from most frequent (“Social support”) to least frequent (“Money”)

| Code | Number of occurrences in sample responses (Part I; n=60) |
| --- | --- |
| Social support | 20 |
| Resource availability | 20 |
| Intrinsic motivation | 13 |
| Structural support | 13 |
| Having knowledge | 8 |
| Other/Vague | 6 |
| Prior success with OS | 4 |
| Money | 4 |

“Abundance of open-source software, preprint servers, sci-hub (what a gem this is!), github, abundance of data repositories, support from colleagues.”

### Which OS aspects have practitioners encountered in their own formal education?

In terms of engagement in OS practices and principles in learning, about half of the surveyed colloquium participants (N=29) stated that they had encountered none such practices in their own formal education (Supporting Information S3-6 Figures). Participants reported engagement with OS practices in their formal education with the following occurrences: *read open access literature* (N=21), followed by *used open code* (N=19), *used open data* (N=17), *shared open data* (N=15), *published results or papers openly* (N=15), *shared own code openly* (N=13), *outreach/science communication* (N=13), *been taught principles of research reproducibility* (N=12), *used open-access online interactive learning resources* (N=11), *been taught principles of research transparency* (N=7), and *open peer review* (N=4).

We predicted that those colloquium participants that had encountered open science-related aspects in their own education were more likely to engage in those aspects. For these predictions, we found evidence for participants using (*P* = 0.027) and sharing open code (*P* = 0.023), in addition to using open educational tools (*P* = 0.012), more frequently when having experienced those activities in their own education. Further, we found evidence for men sharing (*P* = 0.017) and using (*P* =0.028) open data more frequently than women (Supporting Information S6 table).

### How do OS practitioners involved in higher education value OS in teaching and supervision?

In terms of perceived importance of OS practices in research, teaching and supervision, we found strong evidence for such aspects being given less importance in teaching as compared to research (*P <*0.001), but our data did not reveal any such contrast between supervision and research (Fig. 3; Supporting Information S9 Table).

**Figure 3.**
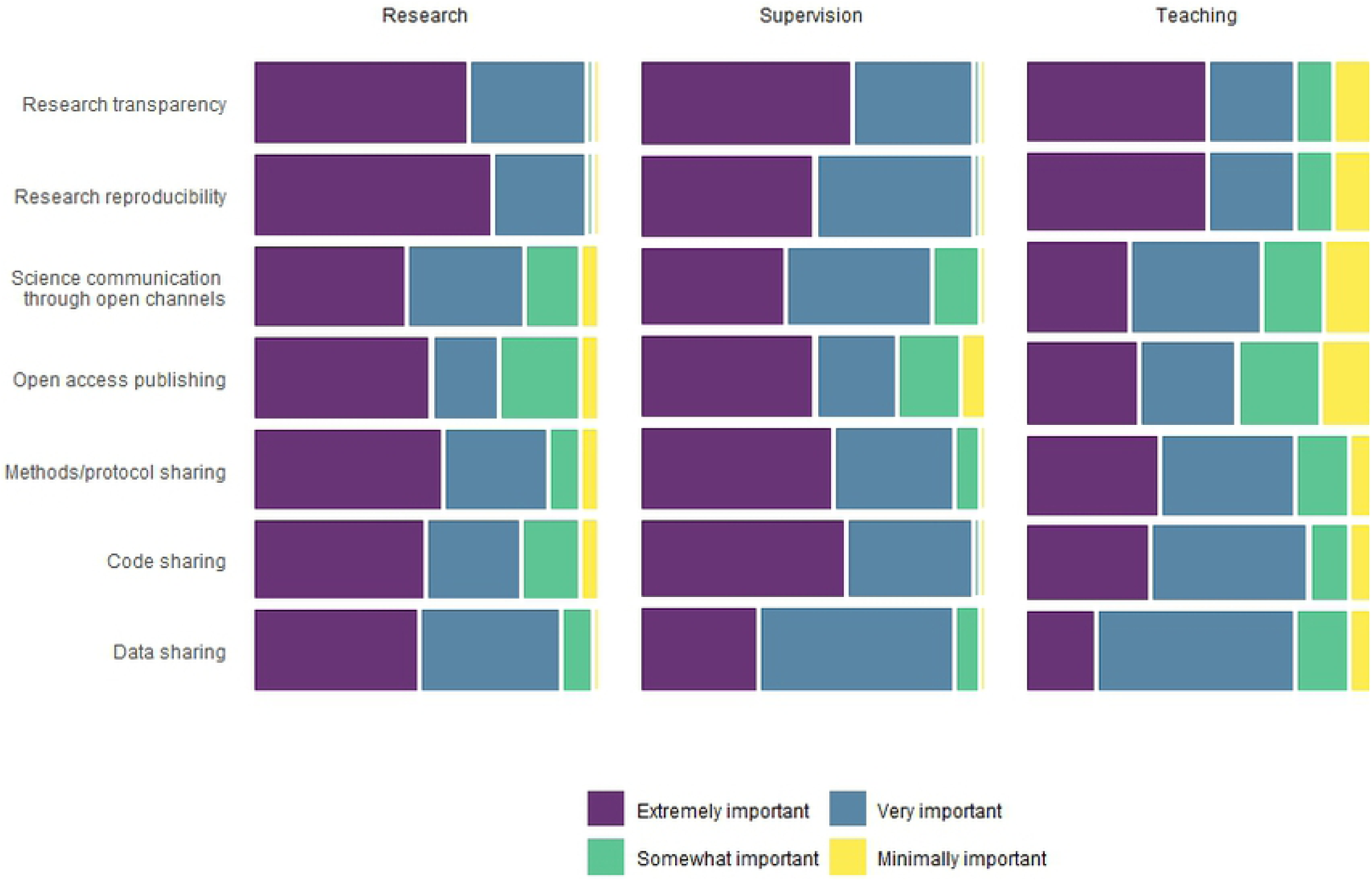
Perceived importance of Open Science aspects for surveyed researchers (N = 26) participating at the colloquium in relation to research, teaching and supervision. Width of tiles corresponds to number of participants involved in the respective activities.

## Discussion

Surveying attendees at the Living Norway 2020 Colloquium gave us a view of how OS can be understood, applied and promoted in a network of dedicated practitioners. We identified two emergent ideas that cut across the responses from the survey, namely 1) that OS is mainly understood in terms of shared and accessible data, code and publications and 2) that individual OS engagement may be further facilitated through education. Firstly, the respondents’ own definitions of OS and the stated frequencies of engagement in related practices revealed that the most emphasised aspects were *sharing data and code openly* in addition to *open access publishing*. Secondly, our analyses indicate that formal instruction may provide incentives to OS engagement as indicated by the participants, while also offsetting disincentives as these are often identified as practical and/or due to lack of knowledge. Still, OS was considered significantly less important in the context of teaching compared to research and supervision. Taken together, these results suggest that an inclusion of OS in teaching and learning can aid in facilitating wide-scale implementations of OS, even though it is clear from our data that this potential may not be evident to educators currently engaging in OS. Thus, a major implication of our study is that by integrating OS principles and practices more formally into higher education, we can naturally address the implementation barriers that depend on individual experience with OS.

Definitions of concepts can reveal how they are perceived and understood, and gathering such definitions is a method that has been applied in previous research (e.g. [14]). Colloquium participants were asked to define OS before the first day of the colloquium, and those definitions helped identify the meaning attributed to OS by attendees ahead of the event. The most frequent codes emerging from those definitions largely mirror practices and principles associated with OS in relevant literature [15], indicating that the respondents were familiar with the associated terms and their meanings. Although there were less responses to the final part of the survey, distributed after the colloquium, where participants were asked to define OS a second time, we interpret the high consistency between initial and final definitions (Fig. 2) as evidence that the event did not substantially change respondents’ definition of OS. Meanwhile, we argue that the most frequent codes emerging from the definitions of OS illustrate a shared understanding of the concept among respondents.

Taking a closer look on the participants’ definitions of OS, the most frequent codes reflect aspects of OS that have a strong practical relevance in ecology [16]. While the methods and approaches of ecological field research reflect the complex variability in the science of ecology and in nature, it is both possible and critically important to ensure the epistemological and computational reproducibility by adhering to the FAIR principles, namely that metohds, protocols and data should be findable, accessible, interoperable and reusable. FAIR was explicitly mentioned in definitions of 11 participants and two of these principles were frequently mentioned separately: *accessibility* (N=38) and – in less explicit terms – *replication/reproducibility* (N=19). This frequent understanding of OS may not be representative of the ecological research community at largem, however, as the survey was carried out amongst attendants associated with Living Norway, a collaborative structure that promotes FAIR principles in ecology and OS in general terms.

Collaborative grassroot structures similar to Living Norway have emerged in recent years [17]. Novel programming tools and enhanced computational power better enable ecologists to address high-level complexity in nature and to generate data across studies and systems. As such methods are highly data-intensive, they require improved alignment of data documentation, management and access across research collaborations and institutions, yielding bigger and more complex datasets. Through analyses of complex systems and research synthesis, ecologists can better inform communities, governments and stakeholders through more accurate predictions that address global concerns, such as ecosystem change [17,18] and functional relationships between environments and organisms [19]. Thus, the wide scale enactment of FAIR principles in ecological research is a means to build robust datasets and analythical pathways that can be put to wider use in the service of science and society.

Grassroot-driven implementations of OS depend on individual researchers adopting these practices, and therefore we asked the colloquium attendees both what hinders and helps their individual engagement in OS. For the perceived hindrances, respondents most frequently identified *knowledge, time* and *lack of guidelines* as the main disincentives. Further, *social support* and *resource availability* were frequently mentioned in relation to what helps individual OS engagement. Taken together, these findings suggest that researchers are more inclined to engage in OS if they have some experience with the practices and if facilitated by their formal and informal working environment both socially and in terms of structural or institutional incentives. Although we cannot assume that this mirrors a wider tendency for the OS movement, we argue that higher education can address both incentives and disincentives to OS engagement, most importantly through facilitating learning of OS tools, principles and practices, offering a OS supportive social environment, and providing structural OS support.

Even though we suggest a potential role for higher education in facilitating OS, an interesting observation from our data is that this possibility may not necessarily be evident to OS practitioners, even not those engaged in teaching. Considering respondents’ perceptions of the importance of OS in research, as well as the more frequent engagement with OS for those having a university affiliation, it is remarkable that such practices and principles were deemed less relevant in educational settings. The reasons behind this discrepancy are probably complex and may involve lack of a tradition in higher education for both teaching and learning the relatively new practices and principles of OS, as well as a lack of learning materials and associated uncertainty on how to incorporate OS in teaching and learning activities [20,21]. Furthermore, OS may be perceived by instructors as more relevant for more advanced students, such as those engaged in thesis work, as the respondents gave similar scores for perceived importance of OS in supervision and research. We suggest that a wider adoption of OS in undergraduate teaching could significantly leverage student engagement and learning [22], and thus speed up the implementation of OS in the wider community. Guidance for undergraduate research and inquiry is now emerging, for example Healey & Jenkins [23] provide recommendations and several examples of such efforts in higher education.

Given the increasing impact of OS on the wider practices and principles of science, including applied science and science communication, early engagement in OS may prove highly beneficial to a growing number of students along the lines described for early career researchers (ECRs) [24–26]. Through the careful inclusion of OS practices in higher education study programmes, educators can offer students a range of activities that increase familiarity with OS and its impact on science itself and the science-society interface, while strengthening the acquisition of domain-specific content knowledge. This is likely to promote students’ future carreers not only in research *per se*, but also in other professions that are informed by or interact with research, such as natural resource management, climate science, medicine, engineering and the science-policy interface, more generally.

Efforts aimed at implementing OS practices in academic institutions involve a variety of agents acting at different levels, namely practitioners (grassroots), institutions (meso level), or political regulation (top-down). While grassroot collaborative structures can be found globally, the most substantial institutional and political efforts are seen in Europe. The League of European Research Universities (LERU) recommends universities to “integrate Open Science concepts, thinking, and its practical applications in educational and skills development programmes” [27]. Further, such implementations are likely affected by political influence manifested as initiatives such as the European Open Science Cloud [28] stemming from the European Commission Single Market strategy [7]. While a majority of academic institutions in Europe are aiming for the adoption of OS practices in strategic terms, successful implementations are still limited [29]. We suggest that such challenges can be addressed through an interplay between the agents that are invested in OS across different levels. Since grassroot practitioners involved in research and teaching are on the frontline for implementing OS in academic institutions, political efforts can generate the necessary large-scale incentives and structural support.

As OS is promoted by a variety of agents, the lack of a unifying definition gives room for diverse interpretations or even skepticism towards OS [30,31]. Other barriers to OS engagement can be practical, such as lacking the required skills, or concerns with the trade-offs pertaining to data sharing [32]. For educators intending to introduce OS in teaching and learning, our main advice is to consider successful initiatives that share a similar purpose. As an example, Project EDDIE (Environmental Data-Driven Inquiry and Explorations) engages students in STEM education by applying active learning methods combined with the use of data repositories that follow the FAIR principles [33]. Further, the International Plant Functional Traits Courses offer training in trait-based ecology through a field campaign grounded in FAIR open science practices, including planning and conducting reproducible fieldwork and data management, and experience with publishing data papers [26,34]. Moreover, educators can obtain formal support for implementing OS in teaching and learning provided through workshops, courses and online-tutorials, and Bossu & Heck [35] offer recommendations on the topic.

In conclusion, our study provide insights into how OS can be understood, applied and promoted within a cluster of practitioners. Respondents seemed to understand and practice OS mainly in terms of providing and/or re-using data and code in addition to open access publishing, but are less aware of how OS can support and promote education. Further, statements pertaining to what helps and hinders individual engagement in OS revealed aspects that can be addressed directly through building a collaborative OS science culture, including through higher education and post-graduate training. Even though we can expect variation in terms of experiences and attitudes across the broader ecological and OS communities, we believe that our results are indicative of some trends that deserve closer consideration. In particular, the differential emphasis of OS in research vs. teaching reflects a prolonged schism in academia where these two scholarly activities are typically regulated by dissimilar mechanisms. Therefore, implementing OS holistically in both research and higher education offers a unique opportunity to bring teaching and research closer together, ultimately advancing knowledge and its applications to the most pressing challenges of our time.

